# Mammalian Pabpc4 is non-essential for development, but has roles in growth, post-natal survival and haematopoiesis

**DOI:** 10.1101/2025.11.22.689676

**Authors:** Matthew Brook, Mathias Lorbeer, Jessica P. Scanlon, Lenka Hrabalkova, Boglarka Nagy, Triin Ounapuu, Melina Michael, Rachael C. L. Smith, John J. Henderson, Joao P. Sousa Martins, Sarah E. Howard, Lora Irvine, Nicola K. Gray

**Author notes:** equal contribution (Matthew Brook, Mathias Lorbeer).

## Abstract

Cytoplasmic poly(A)-binding proteins (PABPCs) are multifunctional RNA-binding proteins which play crucial roles in mRNA translation and stability. In mammals, two family members, PABPC1 and PABPC4 appear widely expressed, but the consequences of their loss of function *in vivo* remain unknown, despite PABPC4 being implicated in a variety of human diseases. We address this knowledge gap using a series of *Pabpc4* knock-out mice. Unexpectedly, we reveal that mammalian PABPC4 is not essential for development, contrary to findings in non-mammalian vertebrates. However, its loss affects birth weight, post-natal growth trajectories and survival, although these were not tightly associated. Growth to adulthood was impacted in a sexually dimorphic manner. Viable Pabpc4-deficient mice allowed us to interrogate roles of Pabpc4 that may affect human health. Anaemia represents a large global health burden, and work in a red blood cell (RBC) model suggested a potential role in haemoglobin synthesis. Surprisingly *in vivo*, we find that Pabpc4 loss did not reduce haemoglobin levels but caused microcytic RBCs and altered RBC distribution width. Moreover, conditional genetic approaches established that this was not a red blood cell intrinsic effect. Taken together, this work provides unprecedented insights into the *in vivo* functions of mammalian PABPC4, and caution against inferring mammalian PABPC function from work in cell-based models and/or non-mammalian species. The generated mouse lines also form a valuable resource with which to further investigate the roles of PABPC4 in health and disease.

**Significance Statement:** RNA-binding protein (RBP)-mediated post-transcriptional regulation is essential for life and life-long health. Cytoplasmic poly(A)-binding proteins are family of RBPs that regulate multiple aspects of cytoplasmic mRNA fate. PABPC4 is an understudied family member, genetically associated with a wide range of human diseases. By creating a series of *Pabpc4* knock-out mice, we surprisingly find it is not essential but is required for normal post-natal survival and growth and normal red blood cell development. Our results emphasise the importance of whole organism studies for understanding mammalian PABP function, uncovering differences between function inferred from other vertebrate species and prior cell-based work. The generated mice also provide a valuable resource for exploring its potential broad roles in human disease.

## Introduction

Extensive molecular studies have revealed the importance of the cytoplasmic PABP (PABPC) family in post-transcriptional regulation, through their complex roles in both global and mRNA-specific regulation of mRNA translation and stability. They are best known for their so-called global roles in promoting mRNA translation and controlling the stability of mRNAs when bound to the poly(A) tail (1–3). Many of their mRNA-specific roles such as mRNA-specific translational repression or activation involve binding to sites other than the poly(A) tail, or interacting with mRNA-specific RNA-binding proteins bound to regulatory sites (2, 4, 5). In some species, PABPCs also participate in regulating mRNA quality control or promoting nuclear-cytoplasmic mRNA transport (3, 6). Recent high throughput studies have exposed our incomplete knowledge of their function, for instance by highlighting that their relative importance in regulating different aspects of mRNA fate can be cell type dependent (7, 8).

Despite intensive study of their molecular cellular roles, whole organism metazoan studies of PABPC functions are sparse. Drosophila PABP is required for embryogenesis, oogenesis, spermatogenesis, and body patterning, and affects neurological function (1). *Caenorhabditis* species have two PABPCs, with their mutation affecting gametogenesis, embryonic survival, lifespan and body volume, although the latter effects may be indirect (1). The PABPC family is more complex in vertebrates (1). In addition to the extensively studied PABPC1 (PABP1, often called PABP), vertebrates encode PABPC4 (PABP4, iPABP) and ePABP (ePAB, PABPC1L), which are highly related to PABPC1 in terms of domain organisation and primary sequence. PABPC1 and PABPC4 are likely widely expressed in adult soma based on mRNA distribution in both non-mammalian and mammalian vertebrates (9, 10). In contrast, ePABP expression is restricted to germ cells, being the predominant PABPC protein in oocytes, while also present in early embryos (1). Its mRNA is detectable at lower levels in the male germline (1). Mammals also encode additional germline PABPs that are highly related to PABPC1 but whose expression is restricted to specific spermatogenic stages (11, 12), and other family members that are distinct from PABPC1 in terms of domain organisation and primary sequence (1, 13).

PABPC1, PABPC4 and ePABP are each essential for development in the non-mammalian vertebrate, *Xenopus laevis* (10). Morpholino-mediated knockdown of PABPC1 or ePABP causes 100% embryonic lethality by organogenic stages (10). In contrast, PABPC4 deficiency results in later 100% lethality by premetamorphosis stage 50, predominantly affecting anterior structures, such as the head, eye and digestive system (10). Deficiencies in any of these PABPCs results in reduced global protein synthesis, consistent with each stimulating translation, and their sharing key protein partners (10). However, whilst self-rescue is efficient, cross-PABP rescue is only partially effective suggesting that individual family members have overlapping, but also distinct, molecular functions in post-transcriptional regulation (10).

In mammals, mouse knock-outs are available for germline PABPs. *Epabp^−/−^* females are infertile with a block in oocyte maturation (14), consistent with findings in *Xenopus* (15), but also have earlier folliculogenesis defects (16), a phenotype not addressed in *Xenopus*. Single or combinatorial knockout of testis-specific PABPs do not affect spermatogenesis (17), perhaps due to overlapping expression with Pabpc1 in the male mammalian germline.

In contrast, information on the *in vivo* roles of so-called mammalian “somatic” PABP proteins PABPC1 and PABPC4, is limited. PABPC1 mRNA is widely distributed across tissues and is predicted to be essential in humans (18). Its overexpression in mouse cardiomyocytes results in physiological cardiac hypertrophy, potentially due to a role in stimulus-induced protein synthesis needed to increase cell size (19). Conditional deletion in a mouse model of chronic myeloid leukaemia suppressed CML proliferation and disease progression (20).

No literature on whole organism studies of PABPC4 in mammals is available. *Pabpc4* was identified as an upregulated transcript in activated human T-cells, although Northern blotting showed a wide distribution across human tissues (9). High-throughput genetic and ‘omics approaches associate it with a range of disease states including cancers e.g. (21–25), immune disorders e.g. (26, 27), neurodegeneration e.g. (28), chronic obstructive pulmonary disease and type II diabetes (29) and host response to viral infection e.g. (30–32). Insights into potential roles in mammals also come from a small number of *in cellulo* studies disrupting PABPC4 function, including knock-down in a transformed erythroblast cell line which profoundly reduced haemoglobin levels, suggesting a pivotal role in haemoglobin synthesis within red blood cells (33).

Here, we explore the roles of mammalian PABPC4 in a whole organism mouse study. In contrast to non-mammalian vertebrates, we find that Pabpc4 is not essential for development, as heterozygous crossed progeny exhibit Mendelian ratios at birth. However, birth weight, survival to weaning, and growth trajectories are altered. The availability of viable mice enabled us to explore its potential role in haematopoiesis. We find that loss of Pabpc4 does not alter haemoglobin levels *in vivo*, but unexpectedly results in smaller (microcytic) red blood cells (RBC), an effect we show is RBC extrinsic.

## Results

### Mammalian PABPC4 and PABPC1 are broadly expressed but with distinct expression patterns

Protein expression patterns can aid prediction of potential biological roles. However, there is a paucity of information on PABPC4 protein expression and where this overlaps with PABPC1, with which it shares at least some molecular functions (e.g. (10, 34, 35)). Thus, PABPC4- and PABPC1-specific antibodies were used to probe a panel of adult mouse tissues (Fig. 1A). With the exception of kidney, Pabpc4 was easily detectable in all tissues but at widely varying levels, being highly abundant in pancreas, but at much lower levels in brain, lung and white adipose tissue. Pabpc1 was relatively abundant in some low Pabpc4-expressing tissues (e.g. brain), and less abundant in some tissues with higher levels of Pabpc4 (e.g. heart, muscle). However, tissues such as pancreas, adrenal glands and uterus had relatively high expression of both, whilst neither were abundant in kidney. In some cases, Pabpc4 appeared to resolve into more than one proteoform (Fig 1A). RT-PCR analysis (Fig. S1A) and sequencing confirmed the presence of four splice variants predicted by EMSEMBL, which differ in the intrinsically disordered region between the RRMs and the C-terminal PABC/MLLE domain (Fig. S1A), and encode proteins with predicted molecular weights of 67.8, 69.4, 70.6 and 72.2 kDa. Data-mining of adult GTex human transcriptomic data, showed some commonality of human *PABPC4* mRNA distribution (albeit lacking in isoform expression data) with that of PABPC4 protein in mouse, with high and low levels in pancreas and kidney respectively, but their relative abundance across tissues differed (Fig.1A and S1B). This may reflect species differences and/or differences between *PABPC4* mRNA-abundance and protein levels, potentially resulting from translational control mechanisms which may be conserved with those of PABPC1 (36, 37).

**Figure 1.**
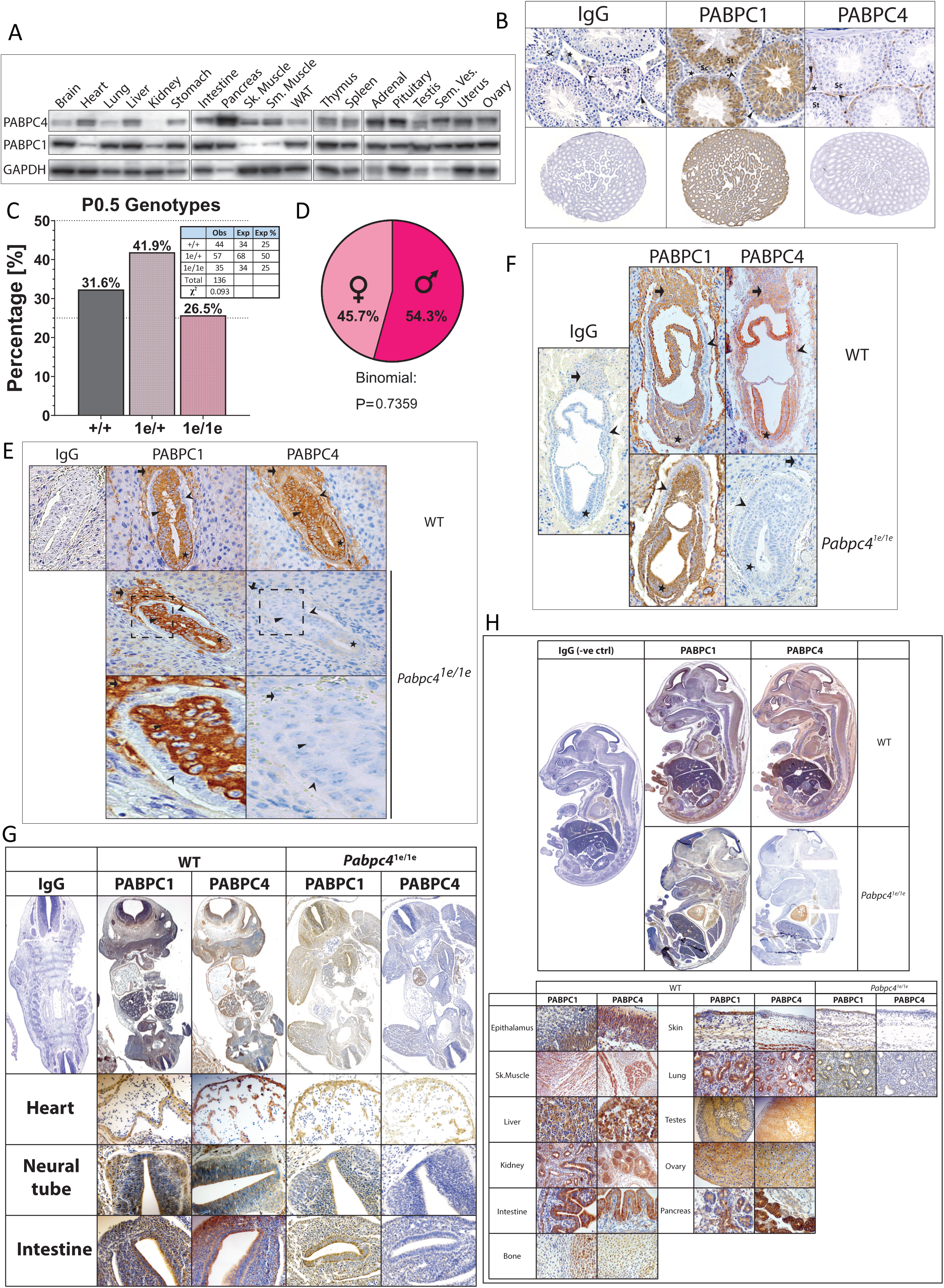
Mammalian PABPC4 is not essential for development. (A) Western blot from indicated adult mouse tissues probed for Pabpc1, Pabpc4 or Gapdh, Sk: skeletal; Sm: smooth; WAT inguinal white adipose tissue (B). IHC with indicated antibodies on murine testis, Star: Leydig cells, chevron: Sertoli cells, arrowhead: spermatogonia Sc: spermatocytes, St: round spermatids. (C). Heterozygous *Pabpc4^+/1e^* cross progeny P0.5 Mendelian ratios and Chi square analysis of deviation from expected values. (D). Sex ratios of *Pabpc4^1e/1e^* mice at P0.5. (E-H) IHC of Pabpc4, Pabpc1 or IgG control in *Pabpc4^+/+^*or *Pabpc4^1e/1e^* embryos and fetuses at e6.5 (E), e7.5 (F), e11.5 (G) and e15.5 (H). Chevron: visceral endoderm; arrowhead: extra embryonic ectoderm, arrow: trophectoderm/ectoplacental cone star: epiblast. Tissue zoom images in (G) and (H) taken from alternate sections where not well represented in main whole embryo sections. *Pabpc4^+/+^*(grey), *Pabpc4^+/1e^* (lilac) and *Pabpc4^1e/1e^* (pink).

To gain insight into relative Pabpc1 and Pabpc4 distribution within adult mouse tissues, immunohistochemistry (IHC) was undertaken (Fig. 1B; and S1C-E). We initially focused on gonads due to prior insight on Pabpc1 immunostaining (17, 38). In ovary (Fig S1C), Pabpc4 was robustly expressed in granulosa cells of different follicle stages, at lower levels in corpora lutea, whereas expression in developing oocytes was more variable, and little staining was observed in the stroma (Fig. S1C). Pabpc1 exhibited a broadly similar pattern but with less oocyte staining, as expected (38), and in keeping with Pabpc1L (ePabp) being the predominant PABPC in oocytes (1). Further confidence in the staining patterns came from the use of previously characterised antibodies, each raised against different PABPC1 or PABPC4 peptides (Fig. S1C, (35), and recapitulation of similar patterns in marmoset ovary (Fig. S1D). Conversely, in testis Pabpc4 and Pabpc1 showed an unexpected reciprocal expression (Fig. 1B), with Pabpc4 being present in mitotic spermatogonia and less prominently in Sertoli cells, whereas Pabpc1 was detectable in meiotic spermatocytes and post-meiotic spermatids, as expected (17). Neither stained the interstitial Leydig cells above background levels. Analysis of non-gonadal tissues, such as cerebellum, also showed non-homogenous expression with striking expression of Pabpc4 and Pabpc1 in the Purkinje cells (Fig. S1E). These data show that individual cell types may robustly express both Pabpc1 and Pabpc4, or predominantly Pabpc1 or Pabpc4, leading us to posit that deleterious phenotypes resulting from loss of Pabpc4 may most likely arise in cells lacking Pabpc1.

### Mouse PABPC4 is not essential for embryonic development

To directly interrogate roles of PABPC4 in mammalian development, a mouse containing a *Pabpc4* gene disrupting insertional mutagenesis cassette (Fig. S1F), *Pabpc4^1e/1e^*, was used. Based on findings in *Xenopus*, we hypothesised that *Pabpc4* would be essential for mammalian embryonic development. Unexpectedly, in mice, all genotypes were represented post-natally (Fig. S1G), and heterozygous crossed progeny were born at Mendelian ratios (postnatal day 0.5, P0.5) (Fig. 1C) with a normal ratio of *Pabpc4^1e/1e^*males to females (Fig. 1D). Thus, these data establish a profound difference in the developmental requirement for PABPC4 in non-mammalian (10) and mammalian vertebrates (Fig. 1C).

Since insertional mutagenesis cassettes do not always completely silence gene expression (39), we sought to determine whether the lack of requirement for PABPC4 for mammalian development may be associated with persistent Pabpc4 expression in *Pabpc4^1e/1e^* fetuses. We also considered whether compensatory changes in Pabpc1 expression, or low or absent Pabpc4 expression during fetal development may play a role. In support of the latter Chorghade et al, 2017 showed that Pabpc1 protein levels are high in fetal heart, but barely detectable in adult heart, raising the possibility that PABPC4 expression may also be different in fetal compared to adult tissue. Thus, Pabpc4 and Pabpc1 expression was examined across development (Fig. 1E-H, and S1H-I), as this has not been determined, in wild-type or *Pabpc4* knock-out embryos. At e6.5, robust Pabpc4 expression was evident in trophectoderm, visceral endoderm, epiblast and extra-embryonic ectoderm, but largely absent from surrounding maternal cells in the implantation site (Fig. 1E). Pabpc1 showed similar expression but was notably absent from the visceral endoderm of the embryo (Fig. 1E). Importantly, given the unexpected viability, Pabpc4 was not detected in *Pabpc4^1e/1e^*embryos, consistent with these mice being a functional null. This also provides additional support for the specificity of the PABPC4 antibody. Moreover, no ectopic compensatory induction of Pabpc1 was observable, most clearly visualised in the visceral endoderm where only Pabpc4 is normally expressed (Fig. 1E). Pabpc4 was also broadly expressed at other developmental stages examined (Fig. 1F-H, Fig. S1H-I), and similar to its expression in adults, it was not homogenously expressed across cell types within tissues (e.g. Fig. 1I, e15.5 skin, lung). Importantly, Pabpc4 was also absent in the *Pabpc4^1e/1e^*embryos at these stages (Fig. 1F-H), with apparent persistent expression in fetal heart being shown to be a cross-reaction with another epitope, by use of a second anti-PABPC4 antibody, (Fig. S1J). Compensatory induction of Pabpc1 expression was also not apparent at these stages (Fig. 1F-H). Thus, mechanistically the lack of an essential requirement for mammalian PABPC4 for development to birth cannot be explained by it only being expressed post-natally, persistent Pabpc4 expression from the *Pabpc4^1e/1e^* allele, or significant compensatory induction of Pabpc1. Combined these data strongly argue that PABPC4 is not essential for embryonic development in mammals, in stark contrast to non-mammalian vertebrates.

### *Pabpc4* is required for normal post-natal growth and survival

To further investigate the biological functions of mammalian PABPC4, we determined whether it played a role in post-natal survival (Fig. 2). Analysis of offspring from heterozygous crosses at weaning (P21) revealed sub-Mendelian ratios of *Pabpc4^1e/1e^* mice, with loss of approximately a third of *Pabpc4^1e/^*^1e^pups (Fig. 2A). Therefore, Pabpc4 plays an important role in survival from birth to weaning. No sex bias was observed in the surviving *Pabpc4^1e/1e^* pups (Fig. S2A).

**Figure 2.**
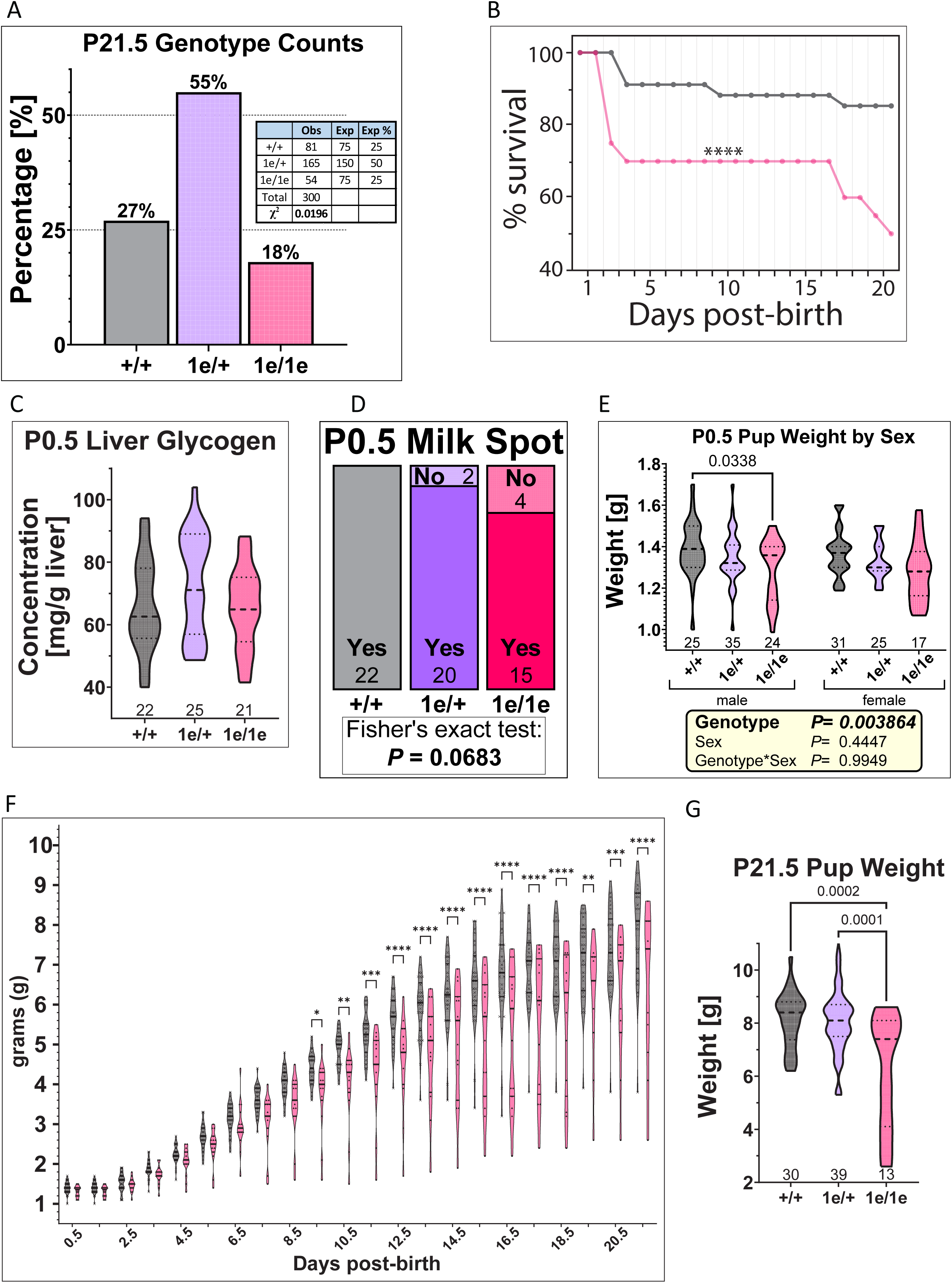
Pabpc4 loss affects post-natal survival and growth. (A). Heterozygous *Pabpc4^+/1e^* cross progeny P21.5 Mendelian ratios and Chi square analysis of deviation from expected values. (B) Survival data of *Pabpc4^+/+^* versus *Pabpc4^1e/1e^* mice weighed daily till weaning. (C-D). Fetal liver glycogen (C) and presence of milk spots (D) are given for all genotypes at P0.5. (E-G) Violin plots of (E) sex-stratified pup weights at P0.5 with tables showing 2-way ANOVA analysis output (F) daily weights till weaning, and (G) P21.5 weight, see also Fig. S2E. Daily *Pabp4^+/1e^* weights are in Fig. S2F. *Pabpc4^+/+^*(grey), *Pabpc4^+/1e^* (lilac) and *Pabpc4^1e/1e^* (pink).

Initially, to gain insight into the mechanistic basis of this phenotype, we determined timing of death and measured birthweight and weight gain in parallel, as growth and survival are often associated. Two main periods of mortality of *Pabpc4^1e/1e^* pups were observed, with most attrition (25%) occurring in the early neonatal period (Fig. 2B). This was borne out by interrogation of breeding colony data (Fig. S2B). Neonatal survival depends on multiple physiological adaptations to *ex-utero* life, and neonatal mortality is therefore a commonly observed phenotype with a poorly understood multifactorial basis (40). Due to the sudden loss of maternal nutrition, survival in the early neonatal period is dependent on pups being born with sufficient nutritional stores and being able to suckle to support their nutrient requirements (41). Thus, measures of nutritional status and feeding; liver glycogen, blood glucose and the presence of milk spots in the stomach were investigated at P0.5, as beyond this time point most deceased pups are routinely cannibalised. This revealed that *Pabpc4^1e/1e^* P0.5 pups have wild-type-like liver glycogen and blood glucose levels, suggesting sufficient available carbohydrate reserves, and most had fed (Fig. 2C-D, and S2C). Importantly, weighing of P0.5 pups revealed that *Pabpc4^1e/1e^* mice had a reduced birthweight (Fig. 2E, genotype effect P=0.0038), a well-recognised risk factor for increased neonatal death (42), providing an attractive mechanistic explanation for some of the increased mortality (Fig. 2B and S2B). Daily weighing of surviving pups also revealed altered growth trajectories and *Pabpc4^1e/1e^*mice did not exhibit catch-up growth and weighed significantly less at weaning on average (Fig. 2F-G, and S2D-F). Most of the deaths after the early neonatal period, in mice weighed daily (Fig. 2B), occurred between P17 and weaning (P21). A broader timing of death was observed in the analysis of colony data (Fig. S2B), consistent with a pleiotropic effect. Some very small *Pabpc4^−/−^* pups were observed (Fig. S2G), and analysis of growth trajectories of individual *Pabpc4^1e/1e^* mice (Fig. S2H) showed that those that died after the early neonatal period, but before wean, initially gained weight but then lost weight prior to death, consistent with a failure to thrive. Interestingly, not all *Pabpc4^1e/1e^* pups born small, or those with reduced growth rates, died (Fig. S2H) with some surviving to weaning, indicating that mechanistically weight and growth rate are not tightly associated with survival in *Pabpc4^1e/1e^* mice (Fig. S2H).

To determine if Pabpc4 deficiency also impacts growth and survival to adulthood, mice were weighed weekly from 4 to 12 weeks of age. *Pabpc4^1e/1e^* mice were smaller at 4 weeks (Fig. S3A), consistent with our data at weaning (P21), and exhibited an altered weight gain trajectory compared to wild-type (Fig. S3B). Intriguingly, sex stratification revealed that this is a sexually dimorphic effect that stemmed largely from the female mice (Fig. 3A), with differences in bodyweight not being statistically significant in male mice (Fig. 3B), although they showed a modestly reduced bodyweight across the timeframe.

**Figure 3.**
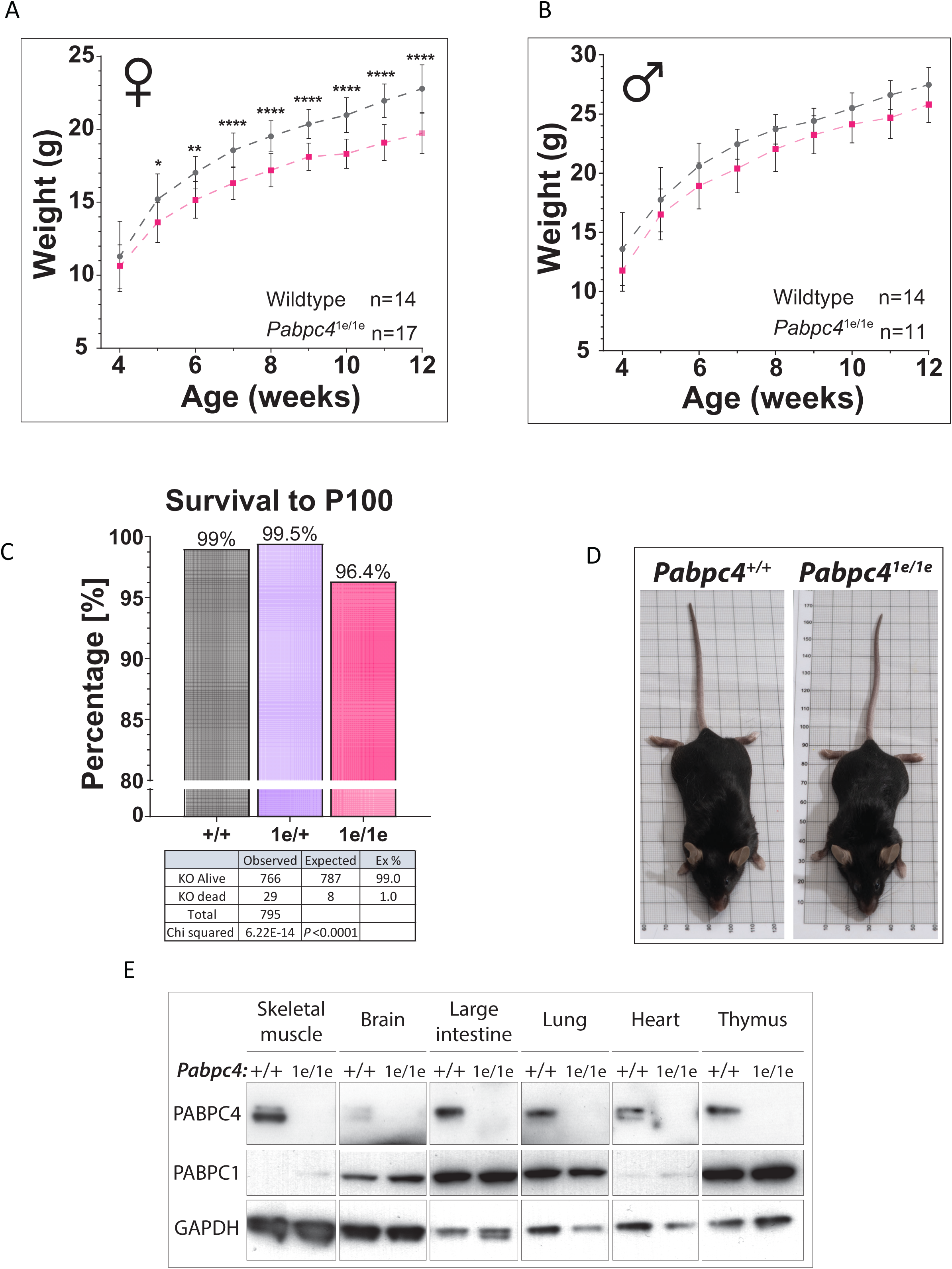
*Pabpc4^1e/1e^*mice can survive to adulthood but show sexually dimorphic differences in growth trajectories. (A-B) 4–12-week growth trajectory for female (A) and male (B) *Pabpc4^+/+^*and *Pabpc4^1e/1e^* mice. (C). Percentage survival of pups of all genotypes from wean to P100, with a split y-axis to allow visualisation of small differences. (D). Photographs female littermates (22 weeks) of indicated genotypes. (E) Western blot of Pabpc4, Pabpc1 and Gapdh in selected *Pabpc4^+/+^* and *Pabpc4^1e/1e^* adult tissues. *Pabpc4^+/+^*(grey), *Pabpc4^+/1e^* (lilac) and *Pabpc4^1e/1e^* (pink).

As none of the weighed weekly *Pabpc4^1e/1e^* mice died, colony data was used to determine if survival from weaning to 100 days was independent of genotype. This revealed a very modest but statistically significant increase in mortality of *Pabpc4^1e/1e^* mice during this timeframe (Fig. 3C; wild-type 99% survival compared to 96.4% for *Pabpc4^1e/1e^* mice). Adult mice had a grossly normal exterior appearance (Fig. 3D) and skeletal X-rays (Fig. S3C), although Scoliosis, or curvature of the spine, and small or missing eyes were noted sporadically, both recognised features in C57BL/6 mice (43); (44). A veterinary pathologist necropsy at 10-weeks did not identify any abnormalities in *Pabpc4^1e/1e^* mice, and organ weights measured as part of this protocol were also found not to be remarkably different in *Pabpc4^1e/1e^* mice (Fig. S3D-H, genotype effect P=0.0877-0.7001), consistent with high levels of post-wean survival. Similar to *Pabpc4^1e/1e^* fetuses, Pabpc4 protein was not detected by Western blotting in a subset of adult *Pabpc4^1e/1e^*tissues, nor was robust induction of Pabpc1 observed, although some tissues showed a small reproducible Pabpc1 increase (Fig. 3E e.g. skeletal muscle). Thus, although Pabpc4 loss impairs survival to adulthood to a very small extent, it is not essential for adult viability. However, it is required for normal growth to adulthood in females. As no other overt morphological or pathological phenotypes were identified in young adult mice, the cause(s) of the reduced growth and survival remain to be elucidated.

### Loss of Pabpc4 results in microcytic red blood cells

The availability of adult mice allowed us to look into potential physiological roles of Pabpc4 that may be represented in human disease. In particular we focused on potential red blood cell (RBC) intrinsic effects on haemoglobin synthesis, as Pabpc4 knockdown in an mouse erythroblast cell line profoundly reduced haemoglobin levels (33), and variants in the human PABPC4-containing locus are genetically associated with multiple RBC parameters, including haemoglobin levels, across multiple studies (45–48). Anaemia represents a major global health burden (49). To address this, we took advantage of a newly developed *Pabpc4* knock-out first conditional-ready mouse (Fig. S1F, tm1a allele; (50)) that allows for deletion of *Pabpc4* in a cell- or tissue-specific manner. Initially, we created whole-body knock-out *Pabpc4^1d/1d^*mice (Fig. S1F tm1d allele), and to our surprise found their haemoglobin concentration and mean cellular haemoglobin concentration were unaltered (Fig. 4A-B; genotype effect P=0.7757 and 0.8848, respectively). *Pabpc4^1d/1d^* also displayed normal RBC number and packed cell volume (Fig. 4C-D). Unexpectedly, both male and female *Pabpc4^1d/1d^* mice displayed a significant decrease in RBC mean corpuscular volume (MCV Fig. 4E; genotype effect P=<0.0001) indicating the RBCs are microcytic. In keeping with this there was also an increase in RBC distribution width (RDW, Fig. 4F; genotype effect P=0.0198), indicating increased variability in RBC size. Thus, Pabpc4 plays a role in RBC development *in vivo*, albeit completely distinct from that previously described in a cell line (33). Our *Pabpc4^1e/1e^* mice, where only females were analysed, also displayed microcytic RBCs (Fig. S4).

**Figure 4.**
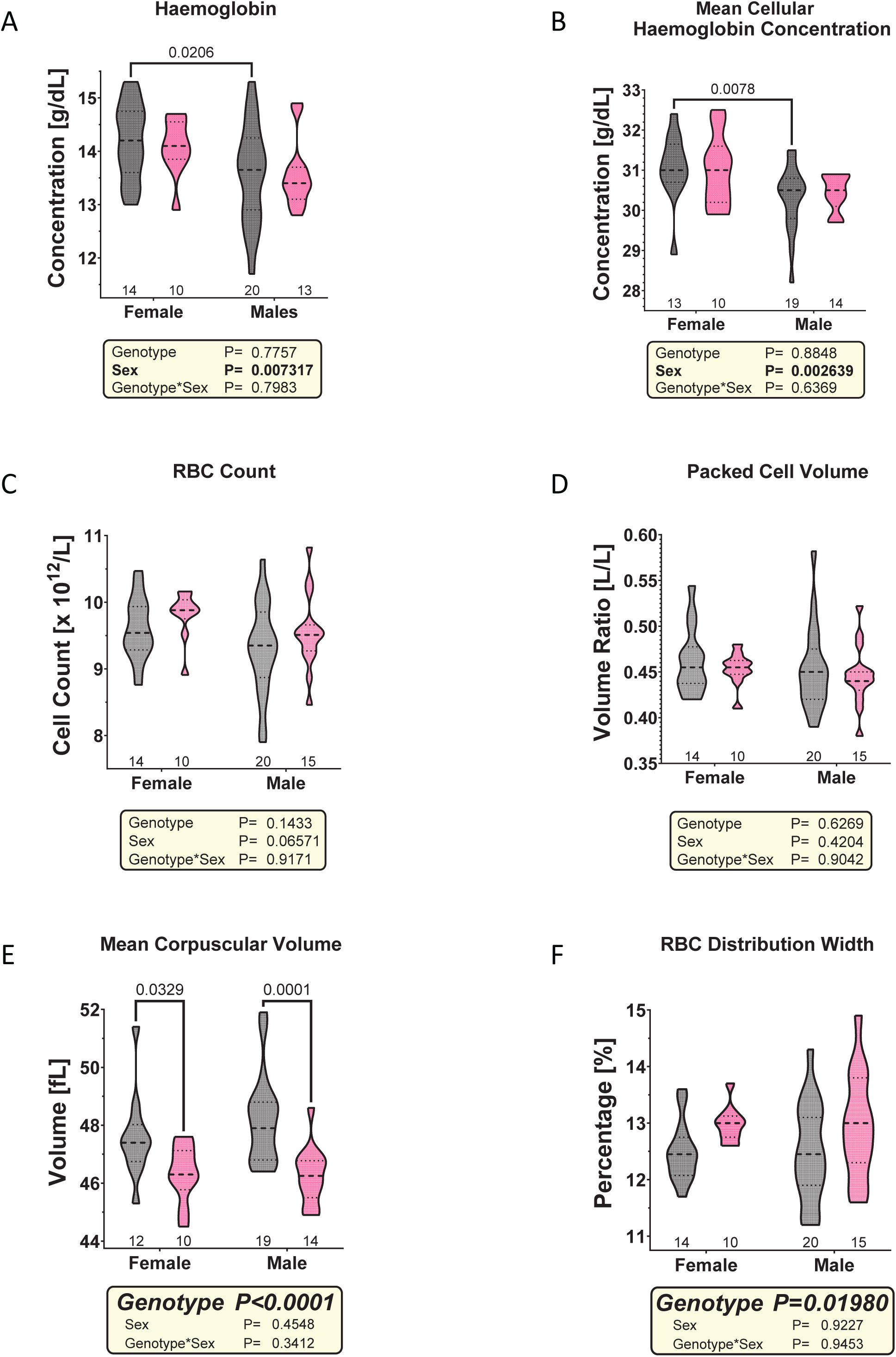
*Pabpc4^1d/1d^* mice have decreased RBC volume and increased RBC distribution width. Violin plots of (A) haemoglobin concentration (B) mean cellular haemoglobin concentration (C) RBC count (D) packed cell volume (E) mean corpuscular volume and (F) percentage RBC distribution width in *Pabpc4^+/+^* (grey) and *Pabpc4^1d/1^*^d^ (pink) mice. Tables show 2-way ANOVA analysis outputs.

To determine whether the mechanism underlying the effects on RBC MCV observed in both *Pabpc4*-deficient mouse lines was due to a requirement for Pabpc4 function within RBCs, a conditional approach was used. *Pabpc4* was specifically deleted within hematopoietic cells by breeding *Pabpc4* conditional ready mice (*Pabpc4^fl/fl^)* (Fig. S1F, tm1c allele) to VAV-Cre mice, a well characterised efficient driver line with few off-target effects (51–53), although it is also efficient in endothelial cells (54). Despite being powered (90%) to detect the RBC MCV difference observed in *Pabpc4^1d/1d^* mice (Fig. 4E), *Pabpc4^fl/fl;Tg(Vav-cre)^* mice did not show a genotype-dependent effect (Fig. 5E; genotype effect P=0.1591), arguing against the altered RBC MCV in *Pabpc4^1e/1e^ and Pabpc4^1d/1d^* mice being RBC intrinsic. Thus, Pabpc4 function within RBCs does not appear pivotal for RBC development as implied from work in cell lines (33), but instead appears to affect their development extrinsically.

**Figure 5.**
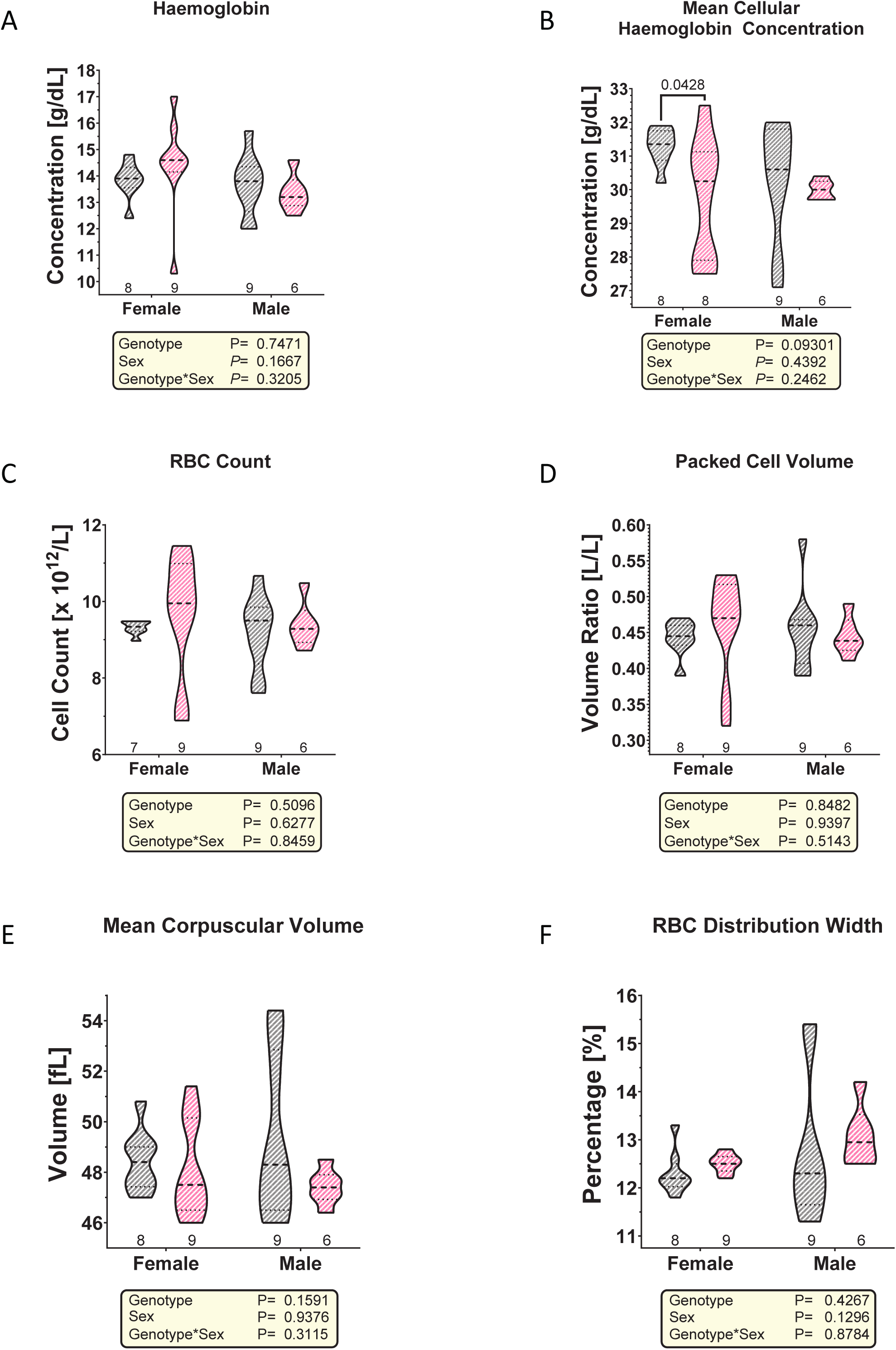
RBC phenotypes are not RBC intrinsic. Violin plots of (A) haemoglobin concentration (B) mean cellular haemoglobin concentration (C) RBC count (D) packed cell volume (E) mean corpuscular volume and (F) percentage RBC distribution width in *Pabpc4^fl/fl^* (light grey) *and Pabpc4^fl/fl;Tg(Vav-cre)^* (light pink) mice. Tables show 2-way ANOVA analysis outputs.

## Discussion

Taken together, our results provide the first insights into the *in vivo* roles of PABPC4 in mammals. Unexpectedly, we find that Pabpc4 is not essential for embryonic development but plays a role in birth weight, and post-natal growth and survival. We also uncover a role in haematopoiesis *in vivo* but which differs in both phenotype and cellular basis from that previously suggested from work in transformed cell lines.

Our data establishes that Pabpc4 and Pabpc1 are widely expressed across and within adult mouse tissues, but in a non-uniform manner, where both can be relatively abundant or low, or where their expression is reciprocal (Figs. 1 and S1). Consistent with their mRNA expression during development in non-mammalian vertebrates (10) we find that Pabpc1 and Pabpc4 proteins are expressed during murine development. Moreover, we uncover that both exhibit similar broad, but non-homogenous, expression patterns during mammalian embryogenesis, with complex patterns akin to our observations in adult tissues (Figs. 1 and S1). Nonetheless our data establish that mammalian PABPC4 is not necessary for basic cell viability, consistent with it not being present in all cells (Figs. 1 and S1). Given the high degree of sequence similarity between mammalian PABPC1 and PABPC4, and the limited studies of mammalian PABPC4 molecular functions showing it also binds poly(A) and AU-rich sequences (33, 34), is polysome associated (55), can stimulate translation (56), and regulate mRNA stability (33, 57), it is possible that a degree of compensation by PABPC1 may contribute to cell viability. However, our results also imply that aspects of mammalian development can proceed without either “somatic” PABP, since neither Pabpc4 nor Pabpc1 are detectable in the critically important visceral endoderm (58) of *Pabpc4^1e/1e^* embryos (Fig. 1). Thus, the idea that cells lacking Pabpc1 may be most sensitive to loss of Pabpc4 is too simplistic, and other factors such as the degree of reliance on post-transcriptional control of different cell types/states likely also affect outcomes.

The presence of multiple Pabpc4 proteoforms (Figs. 1 and S1) adds an extra layer of complexity, as the molecular function of these Pabpc4 proteoforms likely varies due to altered protein and/or RNA interactions. The function of the linker region which is altered by the alternative splicing is less well defined than other PABPC domains but has been implicated in PABPC-PABPC interactions and autoregulation in the case of PABPC1 (59). The specific distribution and relative levels of Pabpc1 and these Pabpc4 proteoforms within different cell types presumably underlies normal tissue function. Splice variants encoding similar human PABPC4 proteoforms strongly supports conserved functional roles (ENSEMBL release 115;(60)). Thus, further study of mammalian PABPC4 molecular functions, including the different proteoforms and the extent to which they overlap with the functions of PABPC1, will also be required to understand the molecular defects underlying different *Pabpc4^−/−^*phenotypes.

Our data unequivocally establish that mammalian PABPC4 is not essential for life (Figs. 1-3). Importantly viable *Pabpc4^1d/1d^* adults containing a deletion in the *Pabpc4* locus were also obtained (Fig. 4), further addressing the possibility that residual non-detectable Pabpc4 expression from the *Pabpc4^1e/1e^*insertional mutagenesis allele explained viability (Figs. 1-3). Viability was unexpected as PABPCs play essential roles in invertebrates and PABPC4 (as well as PABPC1 and ePABP), is essential for non-mammalian vertebrate development (10). Recent work in mouse ES cells, studying LIN28A mediated selective decay of naïve pluripotency mRNAs during the transition to primed pluripotency, an event associated with embryo implantation, suggested a potential requirement for Pabpc1/ Pabpc4 function in mammalian embryogenesis (5). However, the requirement for PABP function was not directly tested in this ES cell study, with work here establishing that Pabpc4 is not essential for *in vivo* embryogenesis in mice (Fig. 1). With regards to species differences, it is interesting to note that PABPC4 had a less broad developmental role and later lethality in *Xenopus* compared to PABPC1 or ePABP (10). Nonetheless, our results caution against directly inferring between species, with conservation of biological roles likely being more nuanced, dependent on family member and phenotype. For instance, ePABP plays parallel roles in oocyte maturation in *Xenopus* and mouse (14, 15), but is only essential for development in *Xenopus* (10). In contrast to our results with PABPC4, this was arguably predictable given the large difference in timing of zygotic transcriptional onset (1 to 2 cell stage in mouse, 4000-8000 cell stage *X. laevis*), which tightly restricts the time period when poly(A)-tail status-regulated gene expression controls mouse developmental programs.

Given the unexpected viability (Figs. 1-3 and S1-3), Pabpc4 and Pabpc1 expression frequently, but not always, overlapping (Fig. 1 and S1), and their sharing of key molecular functions, compensatory changes in Pabpc1 may have been expected. However, robust Pabpc1 induction was not observed when Pabpc4 was absent. Moreover, self- and cross-rescue experiments with Xenopus PABPC1 and PABPC4 show, at least in this species where they share key molecular functions and partners, that they cannot fully compensate for one another, even when expressed using the same regulatory sequences (10), implying they must also have unique molecular functions. A very limited degree of potentially compensatory Pabpc1 induction was observed in very few adult tissues (Fig. 3) but whether this stems from an increase in Pabpc1 expression, or a change in the cellular composition of these tissues remains to be determined. Whilst our results clearly establish Pabpc4 is not essential for mammalian development or survival to adulthood, they do not exclude an essential role for this family in mammals as knock-outs are not available for all family members (e.g. Pabpc1). Nor does the non-essential nature of Pabpc4 for development rule out its contributing to the function of different cell types, that can be revealed with appropriate phenotypic tests e.g. RBCs (Figs. 4-5 and S4). Uncovering the full spectrum of *in vivo* roles of this family will not only require extensive phenotyping of mice harbouring knock-outs of the widely expressed PABPs (Figs. 1, S1), but may also require the generation of compound knock-out mice (e.g. *Pabpc1:Pabpc3:Pabpc6* triple knock-out).

While our study demonstrates the non-essential nature of Pabpc4 for prenatal survival (Fig. 1), the reduced postnatal survival, altered growth trajectories and RBC phenotype clearly highlight important *in vivo* regulatory roles for mammalian PABPC4 (Figs. 2-4). Pabpc4 was required for normal birth weight and post-natal growth, as well as survival to weaning (Figs. 2-3 and S2-3). A particularly high rate of attrition was observed in the first post-natal days (Figs. 2 and S2), and whilst some pups that had not suckled at P0.5 may have gone on to die, this is not sufficient to account for the attrition, nor for those dying after approximately 24 hours (Fig. 3 and S3). Whilst mortality was most prevalent between days P0.5-3.5, it occurred across the whole birth to wean period (Fig 2 and S2) in keeping with a failure-to-thrive phenotype, where pups fail to steadily gain weight, being a multifactorial phenotype. One highly attractive contributory factor in the reduced survival is low birth weight (Fig. 2), a recognised risk factor for post-natal morbidity (42). Consistent with this being a risk factor and not a determinant, not all pups born small, or indeed with low growth trajectories died (Fig. S2). Further work is required to determine whether the reduced fetal growth is an embryo intrinsic or a placental effect, as Fig. 1 shows Pabpc4 expression beyond the embryo proper. Low birth weight and altered growth trajectories are also well recognised risk factors for some adult morbidities (42), and given the observed sexual dimorphism in growth trajectories (Fig. 3 and S3), it is possible that some aspects of adult male and female health may differ in *Pabpc4* knock-out mice.

GWAS studies showed small effect sizes on haemoglobin levels, MCV and RDW (45–48), but follow up studies confirming a role of altered PABPC4 levels in mediating these effects in humans are lacking. Our *in vivo* mouse data did not provide evidence for an association of murine *Pabpc4* with haemoglobin levels, neither was the substantial decrease (approximately 30%) in haemoglobin levels previously seen in a Pabpc4-deficient mouse cell line (33) recapitulated, despite having sufficient power to detect a much smaller difference (Figs. 4-5, and S4). In contrast, our data provide experimental evidence that mouse *Pabpc4* is a determinant of MCV and RDW, which were strongly associated with variants in the human PABPC4-containing locus across multiple ethnicities (P values for MCV: 7x 10^−9^ to 3 x 10^−37^; RDW 2-9 x 10^−44^; 45-48), underscoring the potential relevance of our data to human health. Our results also provide evidence that the human variants act through PABPC4 on MCV and RDW, and thus investigating their impact on PABPC4 levels is now warranted.

Although we cannot rule out a minor RBC intrinsic contribution, our data did not support altered MCV being an RBC intrinsic, or endothelial cell-mediated effect, as the latter are also efficiently targeted by VAV-Cre (Figs. 4-5 and S4). The differences in affected RBC parameters and their cellular basis emphasise the difference between cell-based and whole organism studies of murine Pabpc4. Since the terminal differentiation of red blood cells involves a stepwise destruction of organelles and then RNA content, all of which reduces cell size, the microcytic RBCs in *Pabpc4* knockout mice do not appear indicative of a differentiation block, or high levels of circulating immature reticulocytes (61, 62). As we show that Pabpc4 is widely expressed (Fig. 1 and S1), and reductions in red blood cell size can stem from many factors including nutrient deficiencies such as iron deficiency, auto-immune disease, chronic infection or defects in lipid homeostasis (63–65), further substantive work will be needed to delineate its cellular and molecular basis.

Our study represents the first analysis of *Pabpc4* knock-out mice and form a basis for further studies of clinically relevant phenotypes, including disease states identified in high-throughput or cell-based approaches, phenotypes associated with Pabpc4 partner proteins, or from observations here. For instance, its roles in specific cancers such as prostrate, breast, liver and renal (e.g. (24, 25, 57, 66)) or in host-viral interactions during particular viral infections including coronavirus, herpesvirus and papillomavirus (e.g. (30, 35, 67). Similarly, it remains to be determined whether the effect of Pabpc4 loss on birth weight and growth trajectories (Fig. 2-3) leads to adverse adult metabolic health. Thus, there is still much to be discovered with these different mouse lines providing a valuable mammalian resource in which to tackle such questions.

## Materials and Methods

### Animal breeding

Mice were kept on a 12-hour light/dark schedule, at 20-25°C and 55% humidity with *ad libitum* food and water. All animal procedures were performed under regulation of UK Home Office (Animal Act, 1986) on Project Licences 70-8004 and P4CB1288E and in accordance with ARRIVE guidelines. Knock out first, conditional-ready *Pabpc4* mice were created for us as part of the European conditional mouse mutagenesis program (EUCOMM; www.mousephenotype.org; (68) and obtained from the Sanger Centre (Hinxton, Cambridgeshire, UK) but were later found to be missing the distal loxP site (3’ of exon 3, Fig. S1F) (*Pabpc4^tm1e(KOMP)Wtsi^*[MAUA]). These insertional mutagenesis mice were thus designated *Pabpc4^1e/1e^*and were backcrossed to C57BL6/N. Cohorts in Fig. 3A-B and S3A-B were generated by homozygous breeding to provide sufficient experimental post-wean *Pabp4^1e/1e^* animals. A second knock-out first, conditional-ready *Pabpc4* mouse (*Pabpc4^tm1a(KOMP)Wtsi^*; MEBQ) was subsequently supplied by the Sanger Centre and mated with C57BL/6NTac-Tg(FLPe) mice (69) to create *Pabpc4^tm1c(KOMP)Wtsi^*(conditional-ready) mice (floxed designated *Pabpc4^fl/fl^*) and that were backcrossed to C57BL6/N. *Pabpc4^fl/fl^* mice were mated with B6-Tg(CAG-Cre) mice (70), which removes the critical exon at the one cell stage to give *Pabpc4^tm1d(KOMP)Wtsi^* (whole body null) mice. *Pabpc4^tm1d(KOMP)Wtsi^*knockout mice were named *Pabp4^1d/1d^* and backcrossed to C57BL/6N. MiniMUGA (Transnetyx, USA,(71) confirmed both *Pabpc4^1e/1e^* and *Pabpc4^1d/1d^* mice were C57BL/6N congenic. *Pabpc4^fl/fl^* mice were mated with B6/JOla-Tg(Vav1-cre)1Graf (51) to give *Pabpc4^fl/+;Tg(Vav-cre)^* mice. These were then mated with *Pabpc4^fl/fl^* mice to generate *Pabpc4^fl/fl;Tg(Vav-cre)^* mice in which the critical *Pabpc4* exon was specifically deleted within hematopoietic stem cells and nucleated blood cells.

### Growth trajectories and Survival

New born pups were sexed, genotyped and identified via paw tattooing on day1. Surviving animals were weighed daily at a standardised time of day, to an accuracy of 1 decimal place, and records kept of animals that had died (missing, partial or whole carcasses).

### Necropsy, tissue collection, fixation and gravimetrics

Standardly, animals were culled by Schedule 1 methods: inhalation of rising concentration of carbon dioxide followed by cervical dislocation or cervical dislocation followed by severing a major blood vessel) at 2pm. The following organs were scored for abnormalities during necropsy (Veterinary Pathology Service, University of Edinburgh): heart, lungs, adrenals, ovaries, pituitary, thyroid, uterus, stomach, small intestine, large intestine, mesentery, liver, mammary fat pad, skin, kidney, brown fat, spleen, thymus, lymphosalivary glands complex, pancreas, brain and lower urinary tract (72). Body and organs weights were also determined to 4 decimal points and organs weights were averaged and normalised to bodyweight. Organs were fixed in 4% NBF for 24hrs at room temperature, prior to transfer into 70% ethanol, with the exception of testis and brain. Testis was fixed in Bouin’s solution for 4h then cut in half, fixed for another 2h, before being transferred into ethanol. For brain collections, mice were culled by whole body perfusion with 4% PFA, followed by an additional 24 hours of the isolated brain in 4% PFA, before transfer into PBS.

### Immunohistochemistry

Fixed tissue was embedded in paraffin and sectioned. Following dewaxing, for PABP4 antibodies antigen retrieval was performed in 0.01M Sodium citrate pH 6 (with 0.05% Tween 20 for cerebellum sections), prior to peroxidase inhibition (3% H_2_O_2_ in methanol), water and TBS-T (TBS; 0.1% Tween-20) washes and blocking in 20% horse serum (Biosera Europe, France) with 5% BSA (Sigma Aldrich, UK) in TBS. Immunopurified Rabbit polyclonal anti-PABP1 (395-408, 1:1000) or (623-636, 1:1000) and anti-PABP4 (546-558, 1:1000) or (HPA027301 Atlas antibodies [Sweden], 1 in 1000), were applied in the same blocking solution. IgG (Sigma Aldrich, UK), negative control, was used at matching antibody concentrations. In the case of the alternatively spliced Pabpc4, peptides used to raise the antibodies contain computationally predicted immunogenic amino acids sequences found in each isoform. With the exception of murine ovary and e12.5 embryos, PABPC1 (623-636) and PABPC4 (HPA 027301) were used. After washing twice in TBS-T, an ImmPRESS Horse Anti-Rabbit IgG Polymer (Vector Laboratories, 2BScientific, UK, MP-7401) was applied as per manufacturer’s instructions, prior to two washes in TBS-T. DAB staining was achieved using an ImmPact kit (1drop/ml; Vector Laboratories, 2BScientific, UK), and was stopped by washing in water, before counterstaining with haematoxylin. Slides were mounted using Pertex (CellPath, UK) and imaged using an Axiocam HRc camera mounted on a Zeiss AX10 microscope Olympus Optical, Tokyo, Japan), or a ZEISS Axio Scan Z.1 slide scanner (Zeiss, Oberkochen, Germany) at 20x magnification, using ZEISS ZEN imaging software up to and including version 3.10.

### Blood analysis

Mice were subjected to terminal anaesthesia with isoflurane, and the left ventricle pierced with a sterile syringe to enable blood collection into 1.3 mL EDTA coated microtubes for haematopoietic analysis. Whole blood was analysed by Easter Bush Pathology (Roslin, UK) using an automated flow cytometer-based haematological analyser.

### Glycogen and blood glucose assays, and scoring of milk spots

Neonatal livers were dissected, touch dried and frozen at −80°C. After thawing, 500 µl of 30% KOH saturated with Na_2_SO_4_ was added before incubation at 100°C for 30 minutes. Tubes were flicked to ensure livers were completely dissolved, cooled on ice, and 550 µl 100% EtOH added and shaken well before incubation on ice for 30 minutes to precipitate glycogen, followed by centrifugation at 1,000 g for 30 minutes at 4°C. Supernatant was removed and the pellet dissolved in 1ml of dH_2_O. 25 µl of liver glycogen sample was made up to 1 ml with dH_2_O before addition of 50 µl of 5% phenol in water before 250 µl of 96-98% H_2_SO_4_ was added and were samples mixed and incubated at RT for 10 minutes, before being read at 490 nm. Dilutions of a glycogen standard (Sigma G8876-500MG) was used in parallel to create a standard curve to calculate glycogen concentration. Milk spots were scored following an abdominal excision that allowed for clear visualisation of stomach contents. Blood glucose levels were determined using maltose free Accu-chek strips and an Accu-chek glucose meter (Aviva, Roche) according to manufacturer’s instructions.

### Western blotting

Snap-frozen tissues were homogenised in RIPA buffer containing 2mM DTT, 100µM Sodium Orthovanadate, 50mM Sodium Fluoride, 10mM Sodium Pyrophosphate, 25mM Sodium betaglycerophosphate, 10μM Calyculin A, 1x Complete protease inhibitors (Roche) on ice and extracts clarified by centrifugation at 14000 x g^.-1^. Equal masses of protein were electrophoresed on 4-12% gradient acrylamide gels (Thermo) and transferred to PVDF membrane prior to being probed with anti-PABP1 (623-636) (1:5000), anti-PABP4 (HPA027301, Atlas antibodies, Sweden) (1:10000) or anti-GAPDH (Abcam) antibodies (1:10000). Signals were detected using Suresignal West Femto ECL (Thermo) on a Licor Odyssey Fc imager. Results shown are representative of ≥ n=3.

### RT-PCR

Reverse transcription of Pabpc4 mRNA isoforms was achieved using a First Strand cDNA Synthesis AMV Kit (Roche) with Poly(dT) and random primers according to manufacturer’s instructions, and products amplified by 25 cycles of PCR using Bio-X-Act Long High-Fidelity polymerase (Bioline) using primers I.P4-F1 5’ CAT GCA GAA TTC GCC ATC TTA AAT CAG TTC CAG CCT GC 3’ and I.P4-R1 5’CAT GCA CAT ATG TTA AGA GGT AGC AGC AGC AAC AGT GCC C 3’ that flanked the proposed alternatively spliced region and which contained EcoRI and NdeI sites respectively to allow cloning into pGEM-T-easy and subsequent sequencing.

### Statistics

GraphPad Prism 9 or 10 was used for outlier identification, normality testing and subsequent statistical analysis. Statistical significance of genotype effects was determined by one-way ANOVA with the exception of viability analysis, sex ratios and presence of milk spots which were determined by Chi-Square, Binomial, and Fischer’s exact tests, respectively. Combined sex and genotype effects were determined by two-way ANOVA. The median, 25th and 75th percentile and P values p < 0.05, which was considered statistically significant, are indicated on violin plots shown. Where used one, two or three asterisks denote p < 0.05, p < 0.005 and p < 0.0005, respectively.

## Competing Interests Statement

The authors have no competing interests or disclosures.

## Acknowledgements

We thank Hannah Burgess, William Richardson (deceased) for foundational work, the animal and histology facilities for valuable assistance, Rona Strawbridge for interpretation of GWAS data, Hamish Fraser (previously MRC Human Reproductive Sciences Unit, Edinburgh) for the gift of Marmoset tissue, and past and present members of the lab for useful discussions. Work was funded by MRC studentships (MR/K500963/1, MR/P502030/1) to NKG, a Edinburgh University College Studentship (308406), an MRC program grant MR/J003069/1 and BBSRC project grants BB/R004668/1, BB/V016911/1 to NKG, and an MRC Centre grant MR/N022556/1.

## Author Contributions

<u>Conceptualisation</u>: MB, NKG; <u>Formal analysis</u>: MB, ML, NKG; <u>Funding acquisition</u>: NKG, MB; <u>Investigation</u>: MB, ML, JPS, LH, BN, TO, MM, RCLS, JJH, JPSM, SEH, LI, NKG; <u>Methodology</u>: MB, ML, NKG; <u>Project administration</u>: MB, NKG; <u>Supervision</u>: MB, NKG; <u>Visualization</u>: MB, ML, NKG; <u>Writing-original draft</u>: NKG; <u>Writing – review and edit</u>: MB, ML, JPS, LH, BN, TO, MM, RCLS, JJH, JPSM, SEH, LI, NKG

---

**Figure S1 Expression analysis of Pabpc4 and alleles used** (A) Upper panel: Exonic structure of mouse *Pabpc4* gene and predicted splice variants *Pabpc4* 201-204. Coloured non-coding exons represent alternate transcription start site use (exon 1) and alternate 3’ exon termination (exon 15). Inset shows RT-PCR detection of these splice variants from testis RNA. Lower panel: CLUSTAL alignment of Pabpc4 C-terminal region showing amino acid sequences of the different proteoforms. Amino acid number (RHS) corresponds to that in full length protein. Green, pink, red and blue font indicate the variable linker region, proteoform differences, final amino acids of RRM4 and the PABC domain, respectively (B) Tissue survey of human *PABPC4* mRNA relative expression. GTEx normalised RNAseq quantification (data accurate as of 6/10/2024) (C-E). IHC with indicated antibodies on (C) murine ovary, stars indicated different stages of follicles that contain arrowhead: oocyte, chevron: granulosa cells. Arrow: corpus luteum (D) marmoset ovary, as in D (E) murine cerebellum, Star: Purkinje cells. F) Locus diagrams of the *Pabpc4* alleles used in this study. Where depicted, *Pabpc4* exon numbers are given (grey rectangle, except for critical exon 3 in blue). Insertional cassette containing β-galactosidase (LacZ) and neomycin (neo), SA: splice acceptor; pA cleavage and polyadenylation site; FRT: flippase recognition target site (green diamond); LoxP: locus of X-over P1 site (red triangle). G) Photograph of a P1.5 litter, which genotyping revealed contained 2 WT, 2 KO, as well as heterozygous pups. (H-I) IHC of Pabpc4, Pabpc1 or IgG control in *Pabpc4^+/+^* fetuses at e8.5 (H) and e13.5 (I) with higher resolution of indicated features Br: future brain; G: gut; DA: dorsal aorta; H: heart; S: somites; NT: neural tube; VA: vitelline artery; M: mesothelium; LV: left ventricle; RV: right ventricle; At: atrium; IFT: in-flow tube. When features were not well represented in main whole embryo sections, tissue zoom images in (H) and (I) were taken from alternate sections. (J) IHC of *Pabpc4^+/+^ and Pabpc4^1e/1e^ embryos* hearts at e12.5 with different Pabpc4 antibodies or IgG control.

**Figure S2 Growth and survival of *Pabpc4^1e/1e^* mice to weaning** (A) Sex ratios at wean for the crosses in Fig. 2A. (B) Survival from birth (P0.5) to wean (P21.5) from colony records (C) Fetal blood glucose levels at P0.5 (D-E) P0.5 and P21.5 body weights respectively from only the mice in Fig. 2F which were weighed daily (F) Birth to wean growth curves for all genotypes (G) Photograph of a d13.5 litter comprised of two very small *Pabpc4^−/−^* mice and two *Pabpc4^+/+^*mice. (H) Individual growth curves for mice weighed daily between P0.5 and P21.5. Cohort size is too small to infer any effect on Mendelian ratios at birth. *Pabpc4^+/+^*(grey), *Pabpc4^+/1e^* (lilac) and *Pabpc4^1e/1e^* (pink).

**Figure S3 Growth of *Pabpc4^1e/1e^* mice to adulthood and initial survey of adult mice** (A) Week 4 (28 days) non-sex stratified bodyweights for *Pabpc4^+/+^* and *Pabpc4^1e/1e^* mice (B). Non-sex stratified week 4-12 growth trajectories for *Pabpc4^+/+^* and *Pabpc4^1e/1e^* mice (C). X-rays of 12 week old *Pabpc4^+/+^* and *Pabpc4^1e/1e^* mice. (D-H) Violin plots of normalised sex stratified (D) heart, (E) kidney, (F) liver and (G) spleen and (H) thymus weight in 10-week-old necropsied *Pabpc4^+/+^* and *Pabpc4^1e/1e^* mice. *Pabpc4^+/+^* (grey) and *Pabpc4^1d/1^*^d^ (pink) mice.

**Figure S4 Female *Pabpc4^1e/1e^* mice have decreased RBC mean corpuscular volume**. Violin plots of (A) haemoglobin concentration (B) mean cellular haemoglobin concentration (C) RBC count (D) packed cell volume (E) mean corpuscular volume (F) percentage RBC distribution width in *Pabpc4^+/+^* (grey) and *Pabpc4^1e/1e^* (pink) females.

## Notes

### Competing Interest Statement

The authors have declared no competing interest.

### Summary of Updates

Two additional pieces of data have been added and the text modified accordingly.

