## Supplementary figures and images for "Mammalian Pabpc4 is non-essential for development, but has roles in growth, post-natal survival and haematopoiesis"

### FIg S1

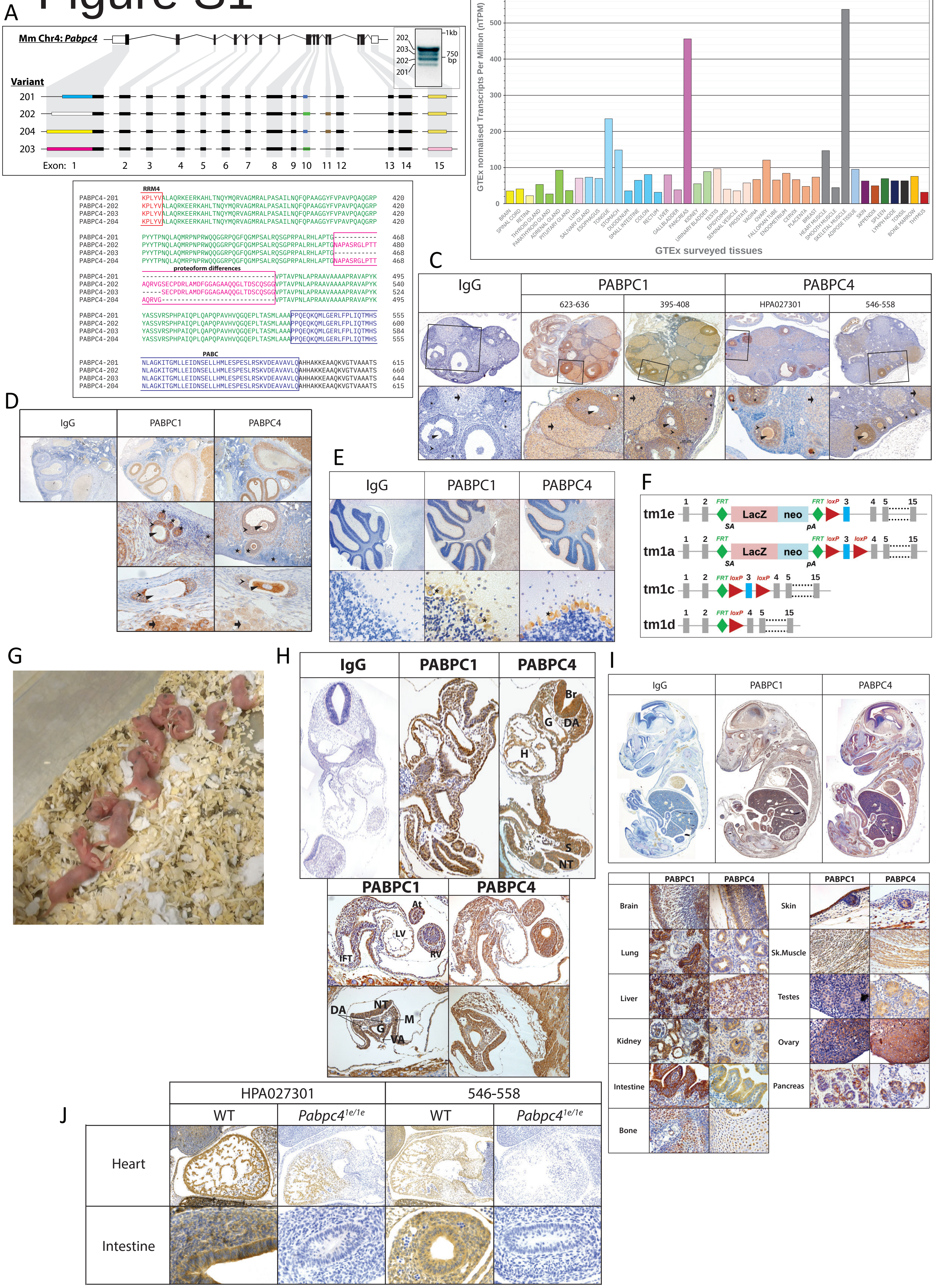

### Fig S2

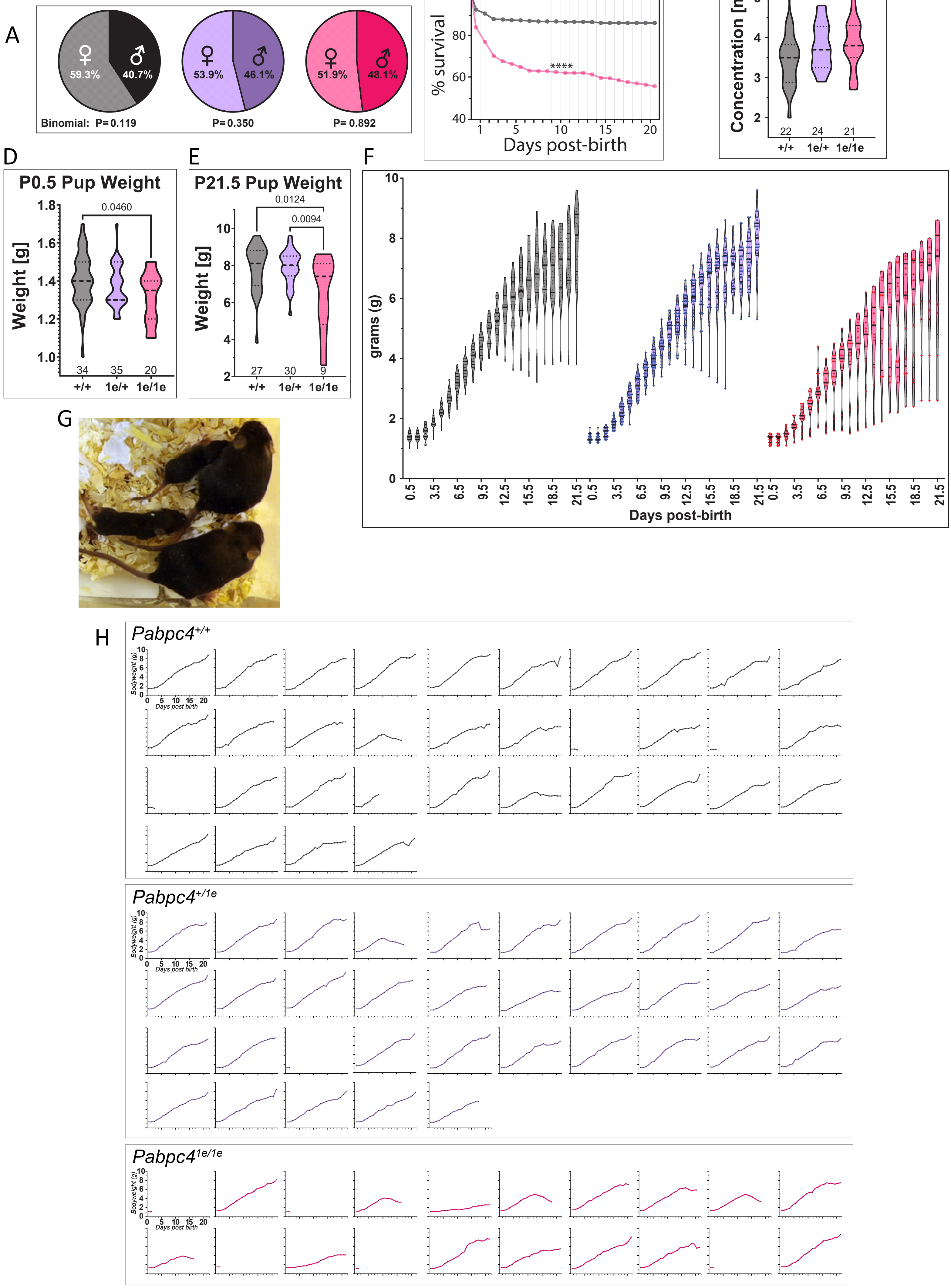

### Fig S3

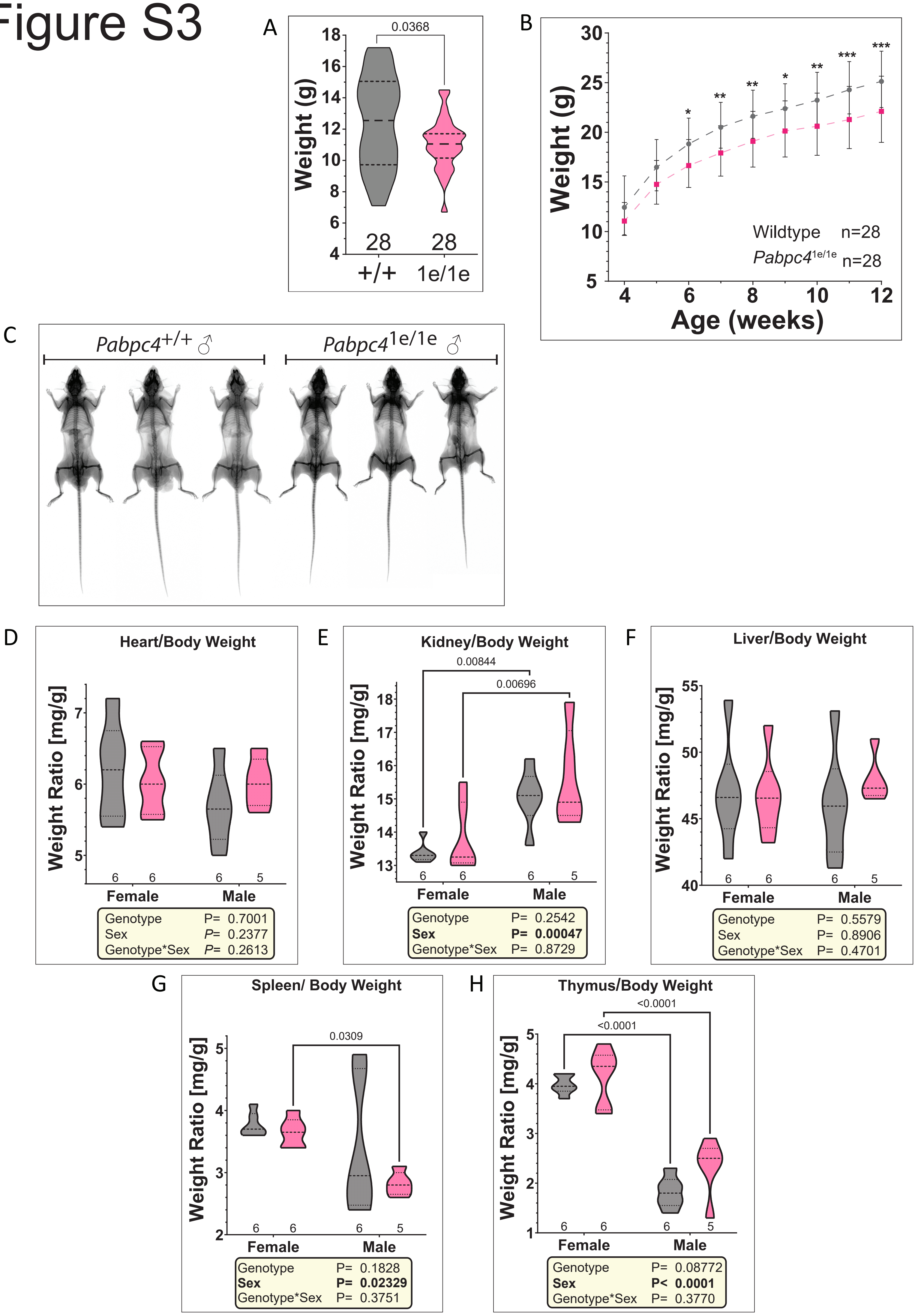

### Fig S4

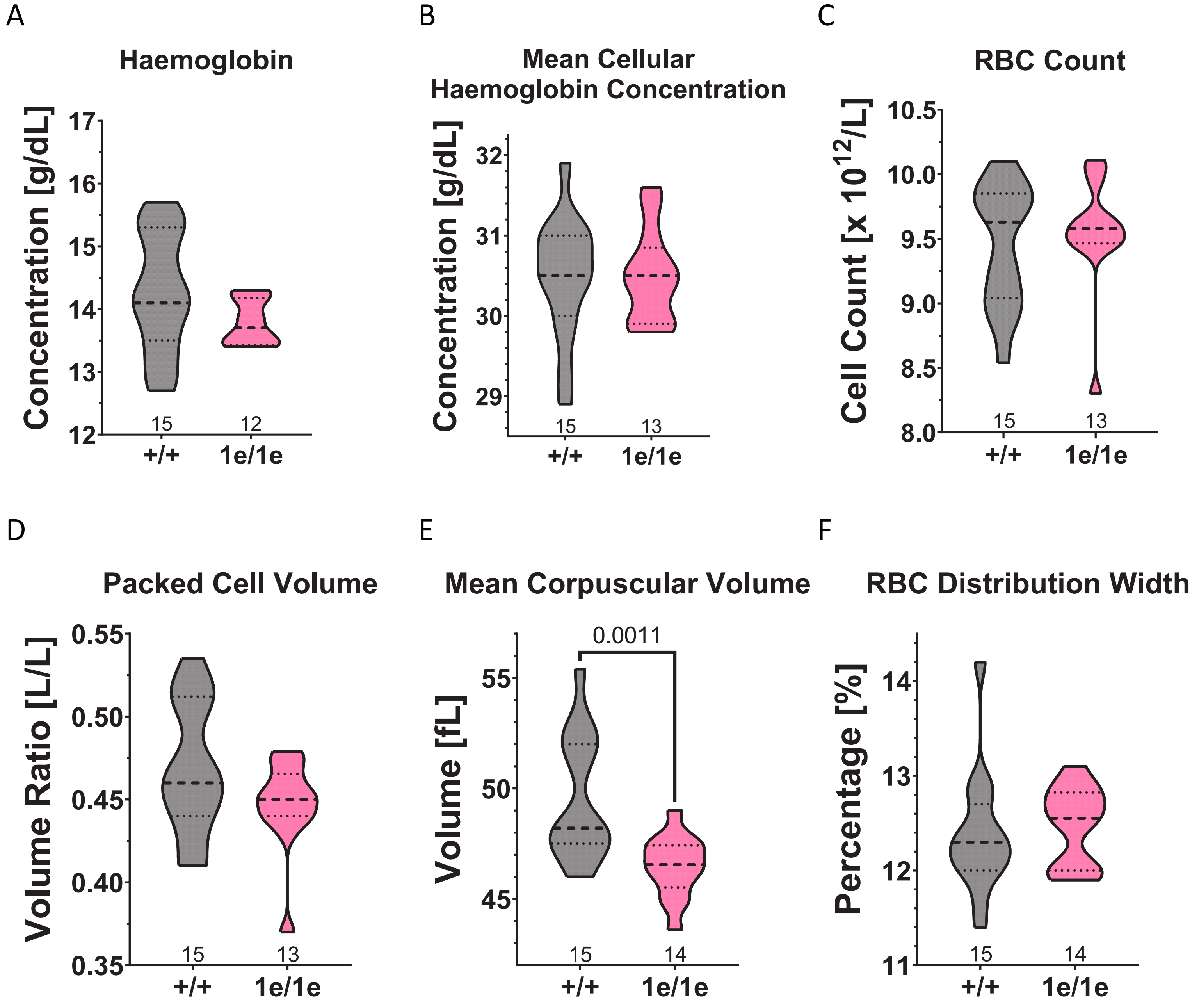
